# Phylogenetic position and expanded host range of the photosymbiont *Brandtodinium nutricula* (Peridiniales, Dinophyceae)

**DOI:** 10.64898/2026.10.26.762032

**Authors:** Nicole L Coots, Leocadio Blanco-Bercial, Gillian H Gile, Patrick J Keeling

## Abstract

*Brandtodinium nutricula* is a dinoflagellate species commonly reported in symbiosis with the marine protist host lineage, Collodaria (Rhizaria). Despite the widespread abundance of this symbiosis throughout the open ocean, and clear indications that the alga should be capable of existing outside of the symbiosis, there are no reports to date of the alga in free-living form in the environment. Even though *B. nutricula* molecular data are publicly available, the phylogenetic position of *B. nutricula* remains unclear. Our taxon-rich 18S rRNA gene phylogenetic analysis revealed a close association between *B. nutricula* and the copepod parasite lineage, *Blastodinium.* We next analyzed Tara Oceans metabarcode data to assess *B. nutricula* abundance outside of the Collodaria symbiosis and found that *B. nutricula* is common in free-living form throughout the sunlit ocean. Similarly, *B. nutricula* does not significantly correlate with Collodaria in terms of read depth in Tara Oceans samples, despite their tight association. We also determined that *B. nutricula* can live symbiotically within planktonic Foraminifera using fluorescence lifetime imaging microscopy in combination with fluorescence in situ hybridization. These findings indicate that the radiolarian-*B. nutricula* symbiosis is less specific than previously suggested, highlighting the complexity of open ocean symbioses and the importance of weighing both host and symbiont perspectives when considering who benefits from the relationship.

## Introduction

Primary productivity in the surface waters of the open ocean has a significant impact on the global carbon cycle and climate regulation. The vast majority of this primary productivity is performed by phytoplankton that are 20 µm or smaller (Detmer and Bathmann 1997; Worden et al. 2004).

Because of their small sizes and relatively nondescript morphologies, studying pico and nano-phytoplankton at the organismal level is particularly difficult. Consequently, basic information about their life histories and interactions are difficult to assess. For example, it was only revealed in the last 15 years that many of these picophytoplankton are mixotrophs, in that they are capable of phagocytosing prey in addition to photosynthesizing (Hartmann et al. 2012; Duhamel et al. 2019). This foundational information dramatically shifted our understanding of how food web dynamics and biogeochemical cycling work in the global ocean (Mitra et al. 2014; Worden et al. 2015; Wilken et al. 2019), thus highlighting the importance and need for basic information about how such an important community of organisms performs daily life functions.

One particularly important alga to the open ocean ecosystem is *Brandtodinium nutricula* (Peridiniales, Dinophyceae). This dinoflagellate was first discovered in association with marine plankton hosts and included within the genus *Zooxanthella* alongside other photosymbiotic dinoflagellates (Brandt 1881). Since then, no one has yet identified *B. nutricula* as a free-living alga in the environment. However, *B. nutricula* is the primary photosymbiont to one of the open ocean’s most abundant host lineages: the Collodaria (Radiolaria, Rhizaria)(de Vargas et al. 2015; Biard et al. 2016). Collodarians are marine protists that form dense colonies that can reach several meters in length (Swanberg and Harbison 1980). Their photosymbioses generate orders of magnitude greater primary productivity than the water surrounding them (Caron et al. 1995), demonstrating the major impact of this relationship on the open ocean ecosystem.

Collodaria are almost never, if ever, observed without *B. nutricula,* indicating a strong functional link between the photosymbiont and the host (Anderson 1976). *Brandtodinium nutricula* actively photosynthesizes within the host, from which the host benefits by assimilating photosynthates and other metabolites (Anderson et al. 1983; Nikitashina et al. 2025). FIB-SEM has revealed that the symbiont changes substantially within the host environment, where plastid volume doubles (Decelle et al. 2021).

These findings suggest host control and host benefit from the symbiosis, but is there benefit to the symbiosis from the symbiont perspective? *Brandtodinium nutricula* has been found as a symbiont to a diverse array of other radiolarian lineages (Probert et al. 2014) and even to the by-the-wind-sailor, a colonial hydroid host (Banaszak et al. 2013). Additionally, *B. nutricula* can be cultured as a free-living alga after physically extracting it from a radiolarian host. While *B. nutricula* loses its flagella and its hard, protective wall (theca) as an endosymbiont, the alga regenerates these traits when removed from the symbiosis and then subsequently grown in culture post-isolation (Probert et al. 2014). Such lifestyle flexibility and host diversity makes the likelihood of a purely exploitative relationship, where the host benefits at the alga’s expense, seem improbable. However, *B. nutricula* has never been documented from the environment in free-living form, probably due to the small size (approximately 10 µm) and unremarkable morphology of *B. nutricula*, which would make identification outside the host environment less likely.

Even though *B. nutricula* is a cultivable alga, surprisingly little information exists about it outside of its role as a symbiont. For example, the phylogenetic position of *B. nutricula* remains contentious. After its original assignment to the genus *Zooxanthella*, *B. nutricula* was later transferred to the genus *Scrippsiella* (Banaszak et al. 1993). Although *Scrippsiella* is occasionally reported as a symbiont to radiolarians (Dolven et al. 2007; Decelle et al. 2012; Liu et al. 2019), the molecular phylogenetics and plate tabulation of *Brandtodinium nutricula* suggested it formed a distinct clade that is not closely related to *Scrippsiella* (Probert et al., 2014). The true placement of *B. nutricula* within the Peridiniales is thus still uncertain. Similarly, all of the ‘omics data that has been generated from *B. nutricula* cultures has only been used for metabolic comparisons between the alga in free-living form versus symbiotic form (Liu et al. 2019; Uwizeye et al. 2021). This molecular data has yet to be used for the purpose of ascertaining *B. nutricula’s* phylogenetic relationships.

Thus, we sought to determine the phylogenetic position of *B. nutricula* using 18S rRNA gene sequencing. We additionally aimed to explore the life of *B. nutricula* outside of the collodarian host. For this, we analyzed global ocean metabarcode data, fluorescence microscopy, and *B. nutricula*’s phylogenetic relationships. Our findings suggest a complex, symbiont-driven lifestyle for an important nano-phytoplankton lineage.

## Methods

### Full-length 18S rRNA gene sequencing

Environmental plankton samples were collected from the open ocean in the Sargasso Sea by dropping a plankton net to 100 m and towing the net vertically to the surface. Individual polycystine radiolarian cells were isolated from environmental plankton samples using a drawn-glass pipette and imaged with DIC optics on an Olympus IX83 inverted microscope. DNA was purified from isolated polycystine cells using the Masterpure DNA & RNA Purification Kit (Epicentre, Madison, Wisconsin, USA). The 18S rRNA gene was amplified from purified DNA using the primers SA-F (Medlin et al. 1988) and V9-R (Sandin et al. 2019), which were initially used to amplify the polycystine radiolarian 18S rRNA gene, but incidentally amplified the dinoflagellate symbiont 18S rRNA gene instead for 6 cells. 18S rRNA gene sequences were ligated into the pCR4-TOPO vector using the TOPO TA Cloning Kit and cloned with One Shot TOP10 chemically competent *E. coli* (Invitrogen, Carlsbad, California, USA) following the manufacturer’s protocols. Inserts from positive transformant colonies were amplified using the sequencing primers M13F and M13R and then purified. All PCR products were sequenced at the Arizona State University Genomics Core Facility on both strands using M13F and M13R primers, plus TAREukF/R as internal sequencing primers (Stoeck et al. 2010).

### 18S rRNA gene phylogenetic tree construction

Peridiniales reference sequences and MALV II outgroup sequences were retrieved from the PR2 database, version 4.14.0 (Guillou et al. 2013). Additional *Blastodinium* sequences and the *B. nutricula* 18S rRNA genes generated and sequenced for this study were added to the fasta file to increase taxon sampling for the Blastodiniaceae. Sequences were then aligned with MAFFT using the L-Ins-I strategy (Katoh et al. 2002) and trimmed with auto settings using trimAl version 3 (Capella-Gutiérrez et al. 2009). Pairwise percent identity scores for Blastodiniaceae sequences were assessed with Geneious v. 2025.1.1. A maximum likelihood TIM2+F+I+G4 18S rRNA gene tree was generated with IQ-TREE version 2.2.2.7 with 1000 ultrafast bootstraps (UFB) (Nguyen et al. 2015). Posterior probabilities were computed with MrBayes version 3.2.7 (Huelsenbeck and Ronquist 2001). Long branches and contaminating sequences were identified by eye and removed from the alignment to prevent long-branch attraction for the final phylogenetic tree iteration.

### Tara Oceans metabarcode datasets

Raw photic zone reads were downloaded from the EBI Sequence Read Archive (Run Accession ERR562 – Metadata for each sample is present in Supplementary Table 1). Sequence reads were denoised with DADA2 (Callahan et al. 2016), and ASVs with a read depth below 10 were removed for quality control. We used BLASTn to identify reads of interest, where the database supplied was a series of Collodaria or *B. nutricula* 18S rRNA gene sequences, and the query was the denoised and filtered ASVs. BLAST hits of 100 bp or longer and with a percent identity of 85 or higher were then added to already-created multiple sequence alignments of Peridiniales or Polycystinea for *B. nutricula* hits and Collodaria hits, respectively, using mafft version 7.505 with the –add flag (Katoh et al., 2002). Maximum-likelihood 18S rRNA gene trees were made with the consequent multiple sequence alignments using IQ-TREE version 2.2.2.7 (Nguyen et al., 2015). ASVs that formed a monophyletic group with exclusively either Collodaria or *B. nutricula* were given the corresponding taxonomic assignment.

*Brandtodinium nutricula* and Collodaria reads were summed according to the size-fraction of the sample from which they originated to determine the relative abundance of both lineages in each size fraction. Linear regression analyses, barplots, scatterplots, and maps of ASV read depth based on sample coordinates were generated in R, where the plots were created with ggplot2 (R Core Team 2013).

### Fluorescence in situ hybridization

Environmental plankton samples were collected by dropping a plankton net to the approximate depth of the Deep Chlorophyll Maximum (DCM) and towing the net vertically aboard the R/V Atlantic Explorer during monthly Bermuda Atlantic Time Series (BATS) expeditions (Supplementary Table 2). Plankton samples were concentrated with gravity filtration through a sieve of the same size as the plankton net. Concentrated plankton samples were fixed in 4% neutral buffered formalin with added 250 µM SrCl_2_ for varying durations. After fixation, plankton samples were washed thrice by allowing the plankton to settle to the bottom of the tube over the course of approximately 1-2 hours, after which point the supernatant was replaced with 0.2 µm-filtered seawater. Fixed, washed samples were stored at 4°C until further processing.

For FISH, we first confirmed that both probes to be used in this study could be used concurrently without competition in terms of the physical location for which they hybridized to the 18S rRNA gene, and that both probes could be used with the same FISH parameters (i.e. formamide content, hybridization temperature, and hybridization duration). For the actual benchwork, we followed a publicly available protocol (Coots et al. 2026): Plankton samples were dried onto Histogrip-coated #1.5 coverslips by incubating the sample on the coverslip in a 46°C incubator for several hours, or until fully dried. Samples were then further dehydrated in a 20 µm HEPES-buffered ethanol series, where the coverslip was incubated in 50%, 75%, and 95% ethanol (twice per concentration) for 3 minutes each. The coverslips were then incubated in the dark in a humidity chamber with a 50 ng/µl probe solution on top of the coverslip (See Supplementary Table 3 for exact reagent solution concentrations). Hybridizations took place for 5 hours at 46°C. After FISH, coverslips were washed in a wash buffer for 30 minutes at 46°C (Supplementary Table 3), then briefly dunked in milliQ water at room temperature.

FISH was performed using two probes concurrently: a *B. nutricula-*specific probe (BNUT) with a Cy5 fluorophore (Coots et al. 2026), and a general eukaryote probe with a Cy3 fluorophore as positive control. After incubation in a wash buffer, coverslips were then stained with DAPI (another control) by incubating the coverslip in 100 µm DAPI in milliQ water at room temperature for 30 minutes and then briefly dunking the coverslip in fresh milliQ water again. Coverslips were then mounted to microscope slides with CitiFluor AF1 mountant solution (Electron Microscopy Sciences, Hatfield, PA, USA) and sealed to the slide with nail polish before imaging.

For each FISH experiment, a control coverslip with dried plankton sample underwent the exact same treatment as the experimental coverslip, except without FISH probes or DAPI staining.

### Imaging & Fluorescence Lifetime Imaging Microscopy

Brightfield and fluorescence imaging took place using the Leica Stellaris 8 FALCON confocal microscope with two multiphoton lasers – one set to 780 nm, and one set to 1045 nm. Three HyD detectors were used to collect fluorescence emissions from each probe, where probes with overlapping fluorescence spectra were imaged in two different channels simultaneously to prevent fluorescence crosstalk. Image acquisition parameters including laser intensity and smart gain were first adjusted to optimize signal-to-noise ratio, then kept constant, until lifetime imaging.

Fluorescence Lifetime Imaging Microscopy (FLIM) was used to distinguish between symbiont autofluorescence and FISH probe signal, using the same method as described in Coots et al (2026). Briefly, plankton samples were scanned for Foraminifera using brightfield microscopy, and each symbiotic foram present was then imaged with each fluorescence spectrum. The fluorescence lifetime for forams possessing brightly fluorescing symbionts in the Cy5 emission spectrum (i.e. the BNUT FISH probe) was collected. If a foram displayed a fluorescence signal in the BNUT FISH probe emission spectrum but not the general eukaryote FISH probe emission spectrum, then the foram was not counted. For lifetime imaging, laser pulsing was fixed at 78 MHz. Laser intensity was adjusted to ensure that photon arrival was below or relatively equal to one photon per laser pulse, and then the FLIM image and data were collected. After image collection, the average lifetime of all photons present within the boundaries of a retarian symbiont region of interest was measured by fitting the fluorescence decay data by applying exponential reconvolution fitting, which is the process of mathematically convolving an n-exponential decay model with the instrument response function (IRF). For this, the number of exponents were iteratively adjusted until the measured decay times had the closest fit to theoretical decay times, and until the chi-squared value for curve-fitting was closest to 1.0. This process yields an average and standard deviation lifetime (in nanoseconds) of all pixels in a region of interest. Here, the intensity-weighted average lifetime and the associated standard deviation were recorded for every symbiont identified (Supplementary Table 4).

Symbiont autofluorescence lifetimes were measured in the same way on the control coverslip. Symbiont fluorescence on experimental coverslips (i.e. those containing FISH probes) was attributed to the FISH probe hybridization when the standard deviation of the symbiont lifetimes on the experimental coverslip did not overlap with the standard deviation of the symbiont lifetimes on the control coverslip (i.e. the lifetimes of symbiont chlorophyll autofluorescence, with no FISH probes applied).

## Results & Discussion

### *Brandtodinium nutricula* is a member of the family Blastodiniaceae

*Brandtodinium nutricula* is sometimes reported as sister to the dinotoms (Kryptoperidiniaceae) (Gottschling and McLean 2013; Cooney et al. 2024), a group of dinoflagellates with photosynthetic diatom endosymbionts (Hehenberger et al. 2014; Yamada et al. 2019, 2024). However, in both studies showing this relationship, taxon sampling for Peridiniales specifically was quite low. According to large subunit and small subunit rRNA gene trees, *B. nutricula* has also been reported as sister to the calcareous dinophytes (Ensiculiferaceae) and *Heterocapsa*, respectively, but with low statistical support (Probert et al. 2014). Lastly, in one analysis based on SSU-ITS1-5.8S-ITS2-LSU concatenated sequences (Li et al. 2020), *B. nutricula* was sister to *Blastodinium,* a genus of marine copepod parasites, but again, with low statistical support (Chatton 1906; Sewell 1951).

To better resolve the phylogenetic position of *B. nutricula*, we performed a taxon-rich 18S rRNA gene analysis focusing on the diversity of Peridiniales (Figure 1). This showed that *B. nutricula* was sister to *Blastodinium* with 97%/0.98 UFB support and Bayesian posterior probability, respectively. Although all sequenced representatives of *B. nutricula* included in this study were isolated and sequenced from the full diversity of known *B. nutricula* hosts, the pairwise percent identity of all *B. nutricula* sequences was 99-100%. This finding supports the assertion that all sequenced *Brandtodinium* symbionts fall within the same species despite a diverse host range. Meanwhile, the pairwise percent identity of *Blastodinium* sequences compared to *B. nutricula* sequences included in this study ranged from 93-97%. The high sequence similarity between *Brandtodinium* and *Blastodinium* sequences raises the question of whether the two should be considered a single genus; however, the significant lifestyle differences and distinct morphologies between the two lineages warrant further investigation into the taxonomy and classification of both. For now, we suggest that *B. nutricula* is a member of the family Blastodiniaceae (Figure 1).

**Figure 1.**
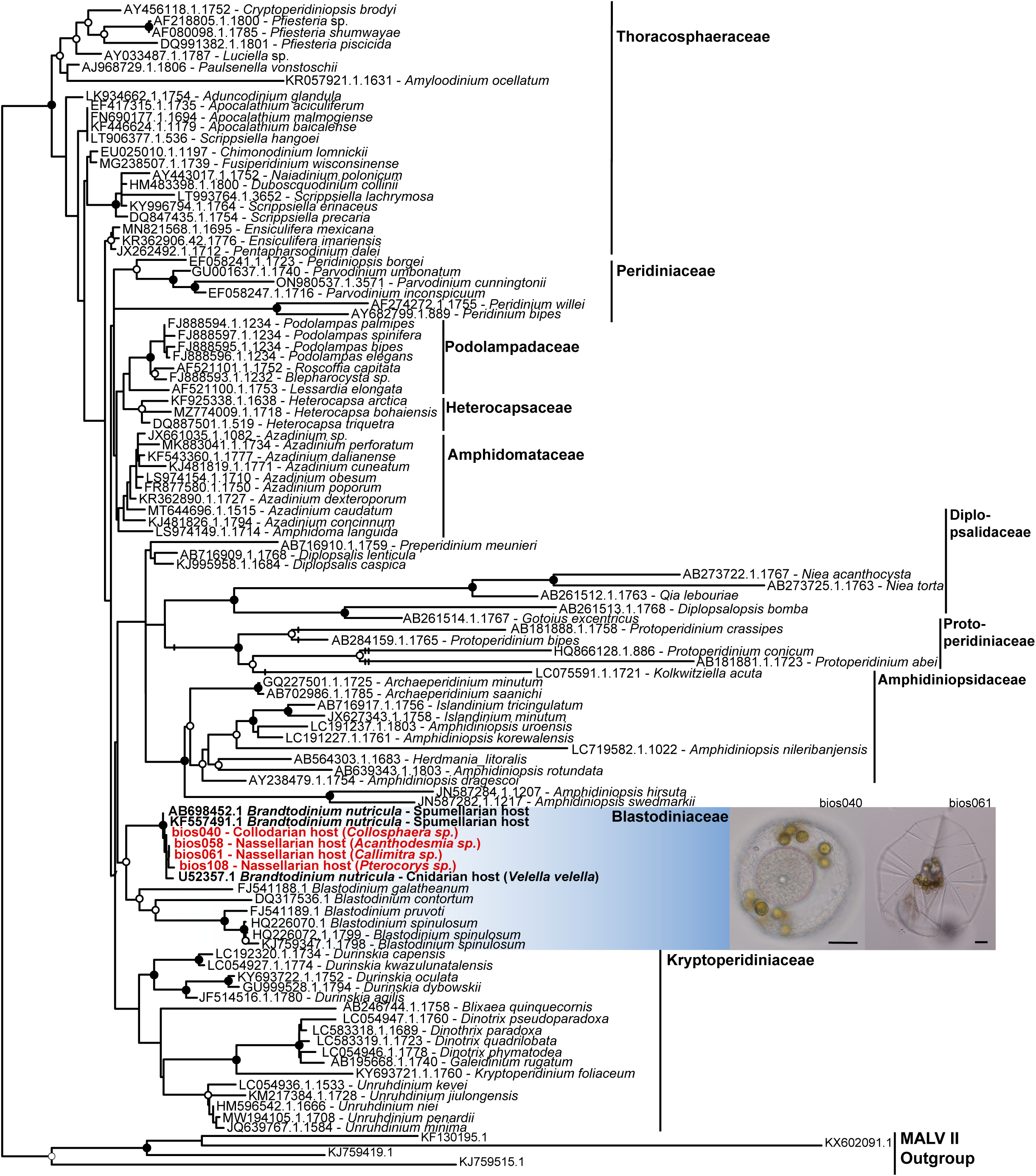
18S rRNA gene maximum likelihood phylogeny of Peridiniales (Dinophyceae). Bold red tips indicate new 18S rRNA gene sequences generated in this study. Branch with a hatch mark indicate that the branch is half of its original length, while double hatches indicate that the branch is one quarter of its original length. Filled circles indicate 100% bootstrap support and 1.0 posterior probability support, while unfilled circles indicate partial support (90-99% bootstrap support and 0.95-0.99 posterior probability support). Images are of isolated radiolarian cells from which two 18S rRNA gene sequences were amplified and sequenced. Scale bars = 20 µm.

This position is interesting given the ecological associations between copepods and radiolarians. Both are highly abundant in the same environments; the surface waters of the oligotrophic open ocean (de Vargas et al. 2015; Fernández de Puelles et al. 2019). Copepods are known to consume radiolarians (Biard 2022), and radiolarians are also known to consume copepods and their eggs via phagotrophy (Anderson et al. 1984; Swanberg and Caron 1991). Many species of *Blastodinium* are photosynthetic, and it has been suggested that their copepod host-association evolved as a means of acquiring nutrients in the nutrient-deplete oligotrophic ecosystem (Skovgaard et al. 2012). This points to a possible evolutionary link as well: one can imagine an ancient ancestor of *Brandtodinium* and *Blastodinium* jumping to a new host via predation in either direction, although the relatively sophisticated and complex parasitic forms in copepods suggest it might be more likely that they are derived from a more general photosymbiosis like what is seen in *Brandtodinium* today.

### *B. nutricula* abundance in global metabarcode data

The global distribution of *B. nutricula* was assessed by determining the abundance of ASVs assigned to *B. nutricula* in photic zone metabarcode samples from the Tara Oceans expedition. Of the 334 photic zone samples analyzed here, 287 possessed *B. nutricula,* with an average read depth of 500+/-1798 (Figure 2A-C, Supplementary Table 1).

**Figure 2.**
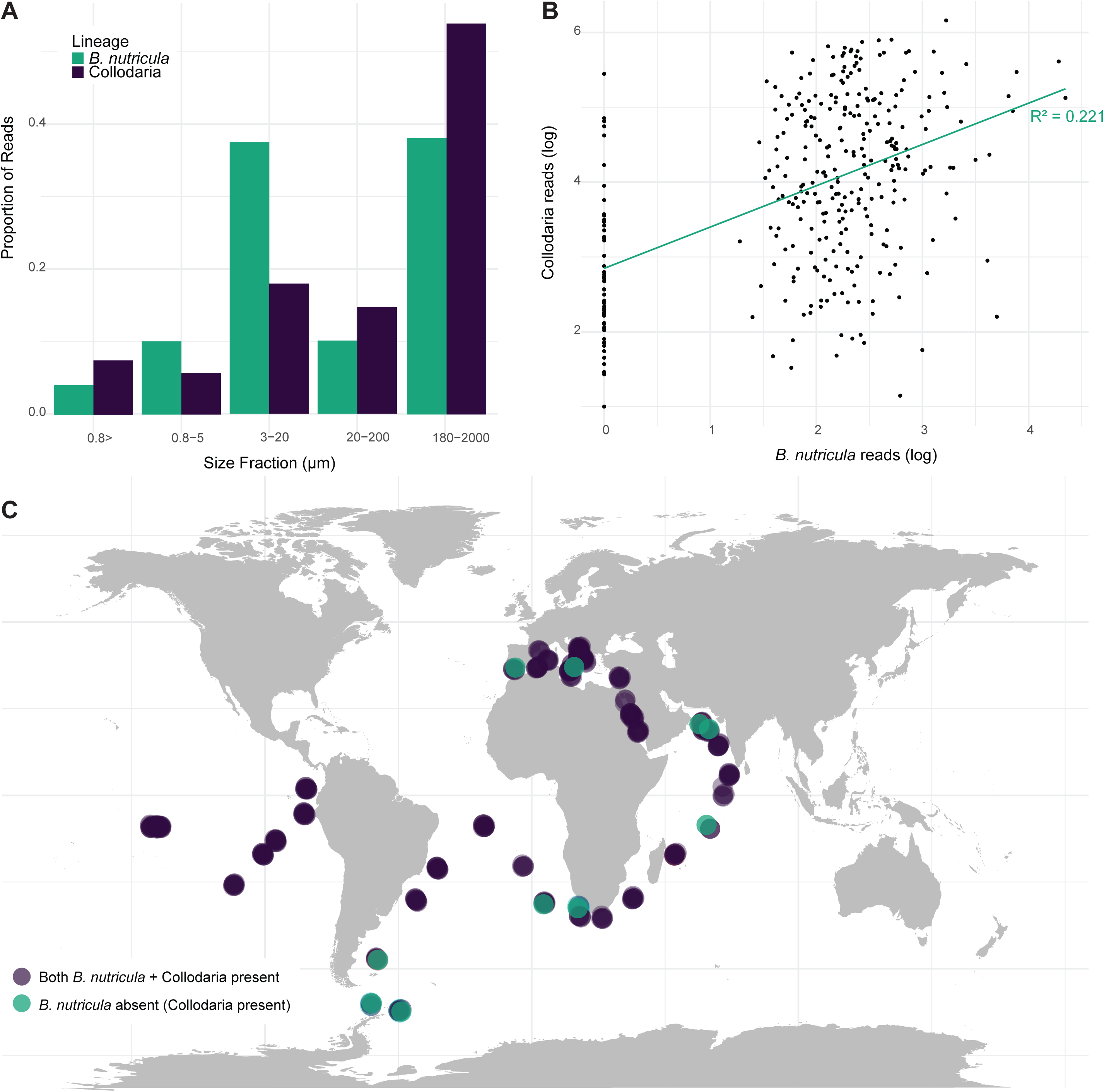
The presence of *B. nutricula* and Collodaria in photic zone samples of the Tara Oceans metabarcode dataset. **A.** The proportion of *B. nutricula* and Collodaria reads in photic zone Tara Oceans samples, grouped by the size fraction from which they were present. **B.** The log-transformed read depth of *B. nutricula* (x-axis) and Collodaria (y-axis) in each of the Tara Oceans photic zone samples, where each point on the scatterplot is one sample. Log transformation of reads was adjusted by 1 such that all samples recorded as zero on the y axis indicate samples in which *B. nutricula* sequences were not detected. The green trendline represents the linear regression of reads across samples, where the R^2^ value = 0.221. **C.** World map portraying the location of all Tara Oceans samples analyzed in this study, and whether or not they possessed *B. nutricula* and/or Collodaria ASVs.

Because collodarians are known to form obligate symbioses with *B. nutricula* alone, we wanted to determine whether there was a correlation between the read depth of Collodaria and the read depth of *B. nutricula* across samples. Collodaria ASVs were present at all 334/334 samples analyzed, where the average read depth of collodarian ASVs across photic zone Tara oceans metabarcode samples was 82,049+/-166,739 (Figure 2C, Supplementary Table 1).

We used linear regression analysis to determine whether there was a correlation in the ASV abundance of collodarians and *B. nutricula* across photic zone Tara Oceans metabarcode samples given that Collodaria are never studied in the absence of *B. nutricula* (Figure 2B). For this, we found that the positive correlation between the ASV abundance of the two lineages was minimal at best (R^2^=0.221, p<2.16E-16) (Figure 2B). This finding is contrary to our expectations because the Collodaria photosymbiosis is known to be highly specific, in which case we would expect a stronger correlation between host and symbiont abundance.

A noteworthy caveat to the linear regression analysis is that 14% of the samples analyzed here did not possess any *B. nutricula* ASVs. Whether or not the absence of *B. nutricula* sequences in some samples represents true absence or artefactual biases remains to be determined. There is no obvious Euclidean or environmental trend that explains why some samples lack *B. nutricula* (Figure 2C); thus, we suspect that *B. nutricula* absence simply reflects sequencing or amplification biases. This would especially make sense given that the relative read depth of *B. nutricula* was quite low across all samples. However, if the absence of *B. nutricula* ASVs in some samples reflects the true absence of *B. nutricula* in those locations, that would suggest that Collodaria can exist without *B. nutricula* more often than previously expected (Biard 2022). Other lineages of Radiolaria display lower specificity to the identity of their photosymbiont associates than Collodaria are thought to display (Decelle et al. 2012; Coots et al. 2025).

Although there is greater evidence to suggest that the Collodaria photosymbiosis is highly specific, one plausible explanation for 14% of Tara Oceans photic zone samples possessing only Collodaria and no *B. nutricula* would be that collodarian photosymbionts have been erroneously identified as *B. nutricula* in the past, and Collodaria is capable of nonspecific photosymbioses like other radiolarians.

Because the linear regression analysis revealed the possibility that host and symbiont abundance vary independently of one another, we next compared *B. nutricula* ASV read depth across size fractions to approximate relative cell abundance in different life stages. Free-living *B. nutricula* should theoretically be present in size fractions of 20 µm or less while the radiolarians that host *B. nutricula* are typically 50 µm or larger (Anderson 1976, 1982; Probert et al. 2014).

*B. nutricula* ASVs were most common in the 180-2000 µm and 3-20 µm size fractions, with 62,928 reads (38.19% of *B. nutricula* reads across the entire dataset) and 61,825 reads (37.52%) respectively (Figure 2A). Meanwhile, 54% of Collodaria reads (or 14,754,651 total) were present in the 180-200 µm size fraction, which represents the abundance of the mature collodarians (Figure 2A). The second-most common size fraction for Collodaria ASVs was the 3-20 µm size fraction, where 17.9% of Collodaria reads (or 49,11,812 total) were present (Figure 2A). This size fraction likely corresponds to the abundance of the swarmer stage of collodarians. Collodarians can also exist as individual entities possessing photosymbionts, where an individual collodarian cell from the Sargasso Sea is typically about 40-60 µm. Individual collodarian cells would therefore fall within the 20-200 µm size fraction, thus explaining the relatively high proportion of reads present for both *B. nutricula* and Collodaria in this size fraction (10.1% and 14.8% respectively) (Figure 2A).

Given that *B. nutricula* is roughly 10 µm in both free-living and symbiotic stages, the nearly identical proportion of *B. nutricula* reads across free-living size fractions (3-20 µm) and symbiotic size fractions (180-2000 µm) suggests that *B. nutricula* could be just as common in free-living form as it is in symbiosis with radiolarians. Still, the method of sample collection could heavily affect the interpretation of relative abundance between size fractions. For example, some radiolarians have delicate skeletons that might be more likely to fracture during plankton net tows, which would lead to symbiotic *B. nutricula* appearing in greater abundance in free-living size fractions if size-fractionation occurs post-tow. The Tara Oceans expedition used a wide variety of sampling methods to collect plankton, however, size fractionation did not occur post-tow for any of the 166 net tow samples analyzed here (Supplementary Table 1). This means that, while it is likely that larger radiolarians and their symbiotic *B. nutricula* could have been present in the smaller size fractions, symbiotic *B. nutricula* are much less likely to be present in small size fractions erroneously.

Assuming that summed sequence read depth in the varying size fractions is an adequate proxy for assessing *B. nutricula* relative abundance, a substantial free-living phase of the *B. nutricula* life cycle would make sense given that radiolarians pick up new symbionts each generation.

While this has never been documented *in situ* due to the nearly impossible nature of studying the radiolarian life cycle, it is known that radiolarian cells divide into hundreds of small, flagellated swarmer cells, approximately 2-5 µm in size (Yuasa and Takahashi 2016). Thus, in the sparse documentation of radiolarian swarmer cells to date, no symbionts have ever been observed, nor is it likely that they could physically possess a symbiont given the difference in size between swarmer cell and symbiont. The small size and aposymbiotic nature of Collodaria swarmer cells also provides biological rationale for our finding that 14% of samples analyzed possessed Collodaria but no *B. nutricula*.

Although horizontal transmission is the most likely scenario for the collodaria photosymbiosis, the relative abundance of *B. nutricula* in free-living size fractions was higher than expected for a photosymbiosis that is thought to be mutually obligate. While horizontally transmitted photosymbioses are relatively common in the marine environment (Russell 2019), the relative abundance of symbiont ASVs in the water column is less than half that of the relative abundance of symbiont ASVs *in hospite* for other obligate symbioses involving horizontal transmission, like the coral-dinoflagellate symbiosis (Lin et al. 2024). Similarly, copepod parasites, which likely have a free-living phase despite an obligate parasitic lifestyle, comprise only 2% of the metabarcode sequences present in marine plankton samples with abundant copepod hosts (Eliassen et al. 2026). These findings support that a symbiosis where the relationship is obligate for the symbiont commonly results in a smaller relative abundance of the symbiont in the surrounding water. Thus, the free-living *B. nutricula* is potentially more common than it would be if the relationship were mutually obligate, so we suspect that *B. nutricula* does not experience the same host specificity to the photosymbiosis as Collodaria does for its symbionts.

The lack of a correlation between host and symbiont ASV read depth, in combination with the finding that *B. nutricula* reads are as common in free-living size fractions as they are in symbiotic size fractions, suggests that *B. nutricula* is highly common outside of the collodarian photosymbiosis, and that the Collodaria-*B. nutricula* photosymbiosis is more flexible in terms of partner identity than previously expected.

### Polycystine radiolarians are not the only symbiotic host to *B. nutricula*

Because *B. nutricula* is related to other symbiotic dinoflagellate lineages, yet their most commonly reported symbiosis is facultative from the symbiont perspective according to metabarcode data, and because *B. nutricula* is already known to form a symbiosis with another large colonial lineage (Banaszak et al. 1993), we sought to determine whether *B. nutricula* can be found in association with other marine planktonic hosts. For this, we applied FISH probes specific to *B. nutricula* to environmental plankton samples collected from the Sargasso Sea.

Doing so revealed a relatively high-frequency association between *Brandtodinium* and planktonic Foraminifera (Figure 3A-H). We scanned plankton samples that were collected throughout the year for the *B. Nutricula*-specific FISH probe and found that 3/17 Foraminifera cells counted from Sargasso Sea possessed *B. nutricula* (Figure 3). Because algal cells often possess substantial, broad-spectrum chlorophyll autofluorescence, we confirmed that the fluorescence signal we counted as our hybridized FISH probe had a significantly different fluorescence lifetime from the chlorophyll autofluorescence of symbionts on a control slide lacking FISH probes (Figure 3B, 3F)(Coots et al. 2026). In both cases where the *B. nutricula* FISH probe was identified within a foram, the total count of *B. nutricula* cells was quite low (only about 1-3 *B. nutricula* cells per foram) (Supplementary Table 4). *B. nutricula* was also only present in the company of other symbiont lineages. This was evident in that the DAPI stain and general eukaryote FISH probe, which were both used concurrently with the *B. nutricula* FISH probe, identified significantly more symbionts present within the cell than could be accounted for by the *B. nutricula* FISH probe alone (Figure 3C-D, 3G-H), and also in that the far-red fluorescence lifetime of other symbionts within the cell was equal to that of chlorophyll autofluorescence (Supplementary Table 4).

**Figure 3.**
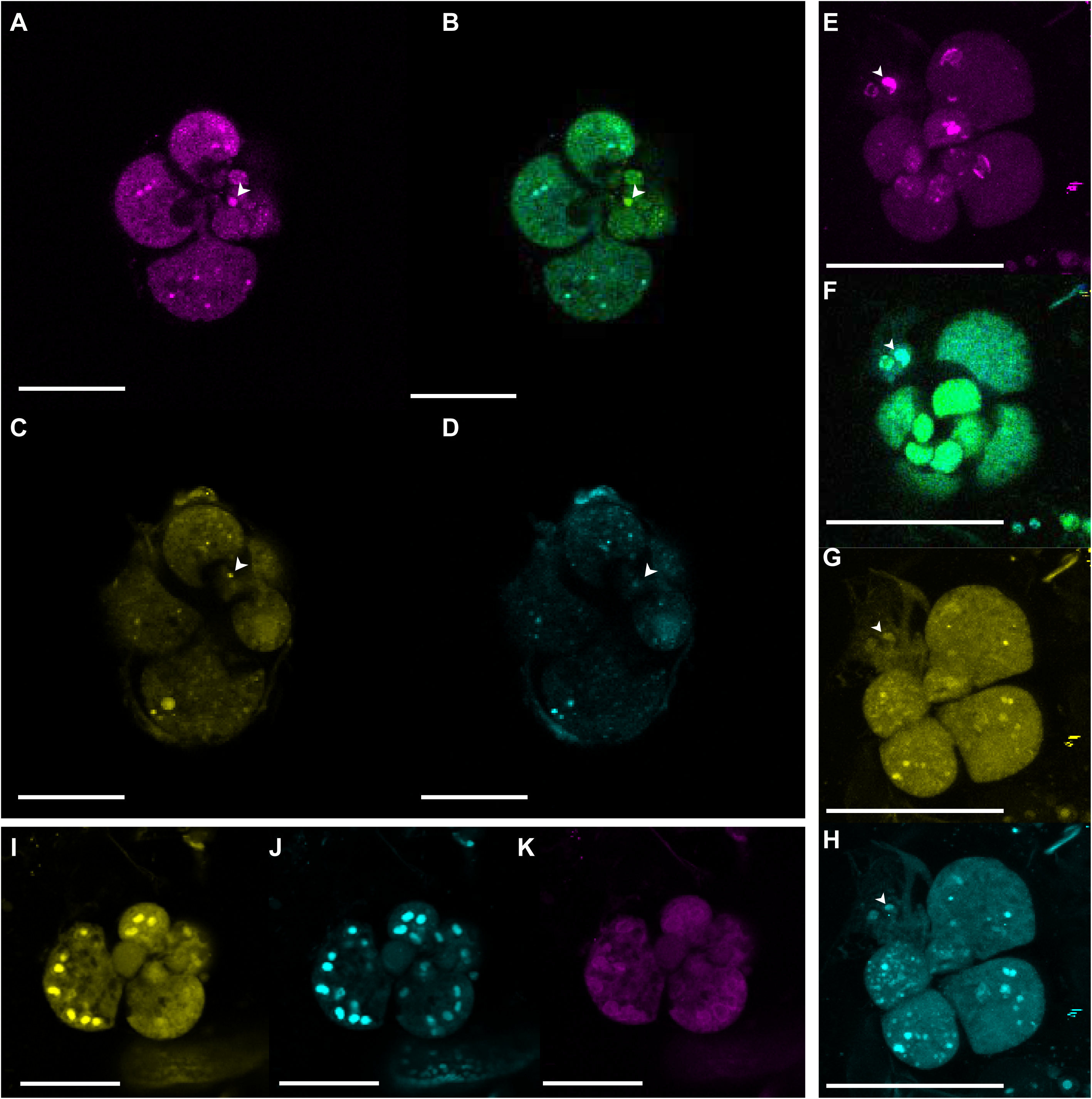
Fluorescence in situ Hybridization microscopy of *B. nutricula* in Foraminifera cells. Scale bars = 25 µm. Colors for each micrograph are the wavelengths imaged as follows: *B. nutricula-*specific FISH probe conjugated with Cy5 fluorophore (far-red/magenta); Heatmap of the fluorescence lifetime of every photon in the far-red wavelength spectrum in the cell (blue to green/rainbow coloration); General eukaryote FISH probe conjugated with Cy3 fluorophore (yellow); DAPI DNA stain (blue). Arrows indicate *B. nutricula* symbionts which were identified based on the significantly different fluorescence lifetime of the symbiont compared to the autofluorescence of the foram and the chlorophyll autofluorescence of foram symbionts on a control coverslip. **A-D.** Foram cell imaged on July 23, 2025, with one *B. nutricula* symbiont identified, among other non-*B. nutricula* symbionts. **A.** *B. nutricula-*specific FISH probe conjugated with Cy5 fluorophore (far-red/magenta) **B.** Heatmap of the fluorescence lifetime of every photon in the far-red wavelength spectrum in the cell (blue to green/rainbow coloration) **C.** General eukaryote FISH probe conjugated with Cy3 fluorophore (yellow). **D.** DAPI DNA stain (blue). **E-H.** Foram cell imaged on November 20, 2025, with one *B. nutricula* symbiont, among other non-*B. nutricula* symbionts **E.** *B. nutricula-*specific FISH probe conjugated with Cy5 fluorophore (far-red/magenta) **F.** Heatmap of the fluorescence lifetime of every photon in the far-red wavelength spectrum in the cell (blue to green/rainbow coloration) **G.** General eukaryote FISH probe conjugated with Cy3 fluorophore (yellow). **H.** DAPI DNA stain (blue). **I-K.** Foram cell imaged on November 22, 2025, with many symbionts, but no symbionts possessing the *B. nutricula* FISH probe. **I.** General eukaryote FISH probe conjugated with Cy3 fluorophore (yellow). **J.** DAPI DNA stain (blue). **K.** symbiont autofluorescence in the far-red wavelength spectrum (magenta).

Of the 17 Foraminifera cells counted, only 3 were aposymbiotic, in that no symbionts were identified at any wavelength. The remaining 14 cells contained between 4-50+ additional nuclei in the cells based on DAPI staining (Figure 3J, 3H). These 14 cells possessed symbionts that yielded a fluorescence signal from hybridization with the general eukaryote probe in addition to DAPI (Figure 3I, 3G), and some additionally possessed chlorophyll autofluorescence, however, the far-red autofluorescence lifetime in those cells was significantly different from that of the FISH probe signal (Supplementary Table 4).

Although Foraminifera photosymbioses are relatively well-studied, surprisingly little information exists about the symbiont identities of the planktonic Foraminifera. They are generally thought to possess dinoflagellate symbionts (Shaked and de Vargas 2006; Siano et al. 2010), however, chlorophyte, chrysophyte, and cyanobacterial photosymbionts are possible as well, and some planktonic Foraminifera can host multiple algal lineages at the same time (Faber et al. 1989; Shaked and de Vargas 2006; Bird et al. 2017; Coots et al. 2026). Our findings suggest yet another new symbiont lineage to planktonic Foraminifera, and yet another new host lineage to *B. nutricula*.

## Conclusion

There are several phytoplankton lineages that commonly exist as photosymbionts to diverse host lineages, with little to no host specificity, like the chlorophyte genera *Tetraselmis* and *Chlorella* (Karakashian and Karakashian 1965; Douglas 1983; Gast and Caron 2001; Androuin et al. 2020; Jenkins 2024). This dynamic suggests that either the symbiont lineage is a generalist that possesses traits that make it easy to be taken up as a symbiont by diverse hosts, or that the symbiont has some control over the establishment of the relationship in diverse hosts. Either scenario would qualify such phytoplankton lineages as “professional symbionts” under our definition of the term. The concept of a professional symbiont, i.e. a symbiont lineages that is adapted to an intracellular lifestyle with little host specificity, is relatively well-documented across several bacterial lineages, including the Rickettsiales and the Holosporales (Hammerschlag 2002; Salje 2021; Castelli and Petroni 2025).

Because *B. nutricula* has a substantial free-living phase, falls within a host-associated dinoflagellate family, and can act as a symbiont to diverse hosts, we suggest that *B. nutricula* is a professional symbiont with high host specificity, but not as restricted as was previously thought. Mixotrophy is a known, common strategy, with clear benefits in the nutrient-deplete open ocean (Selosse et al. 2017; Ward 2019). Given the recent surge in literature suggesting that even the smallest phytoplankton lineages display mixotrophy to some extent, we speculate that professional symbiosis can emerge from similar needs: to give the symbiont access to a stable source of nutrients in an otherwise nutrient-deplete environment. We often focus on the benefit of photosymbiosis to the host, but symbiont-driven photosymbioses may also be an overlooked mode of metabolism for open ocean phytoplankton lineages in cases like this, where the symbiont can readily transform to a free-living stage.

## Acknowledgements

This work was supported by a grant-in-aid from the Bermuda Institute of Ocean Sciences to NLC, a grant from the Gordon and Betty Moore Foundation (https://doi.org/10.37807/GBMF9201) to PJK, and a grant from the US National Science Foundation to GHG (DEB-2045329). We thank the Bermuda Atlantic Time-series Study (NSF OCE 2241455) for providing monthly fixed environmental plankton samples for this analysis. LBB acknowledges the funding from NSF OCE 2227766 and the Simons Foundation International’s BIOS-SCOPE program. The authors also acknowledge the UBC Bioimaging Facility, for which the fluorescence microscopy in this project would not have been possible.

## Data Availability

All 18S rRNA gene sequences produced for this study are publicly available in GenBank under accessions PZ779417-PZ779420.

## Abbreviations

FLIM: Fluorescence Lifetime Imaging Microscopy
FISH: Fluorescence in situ Hybridization

---

**Supplementary Table 1.** Tara Oceans photic zone sample collection metadata and total ASV read depth of *B. nutricula* and Collodaria in each sample

**Supplementary Table 2.** Environmental plankton FISH sample collection metadata

**Supplementary Table 3.** FISH solution reagents and concentrations

**Supplementary Table 4.** FLIM data for all symbiont cells with fluorescence in the Cy5 emission spectrum measured

## Notes

### Competing Interest Statement

The authors have declared no competing interest.

